# Virus infection might cause cells producing black swimming dots

**DOI:** 10.1101/431965

**Authors:** Shuang Tang, Xiao Yang, Xinlong Jiang, Wenzhong An, Xiangchen Li, Shuo Wang, Hongtu Li, Zhengchao Wu, Feng Su, Shiliang Ma

**Affiliations:** Laboratory of Animal Cell and Molecular Biology, College of Bioscience and Biotechnology, Shenyang Agricultural University, Shenyang, Liaoning 110866, China; Instrumental Analysis and Test Center, Shenyang Agricultural University, Shenyang, Liaoning 110866, China; College of Animal Science and Technology, Zhejiang A&F University, Lin’an, Zhejiang 311300, China; Key laboratory of Genetics and Reproduction Health of National Health Commission, Liaoning Province Research Institute of Family Planning, Shenyang, Liaoning 110031, China; College of Animal Science and Veterinary Medicine, Shandong Agricultural University, Taian, Shandong 271018, China

**Keywords:** virus, black swimming dots, cell culture, serum, tissue extracts

## Abstract

Black swimming dots (BSDs) are nanoscale dot-like contaminants in the dishes of cultured cells. Until now, the identity of BSDs has not yet been determined. In our recent study, we proposed that BSDs *per se* are nonliving inorganic nanoparticles yet should derive from the cells infected with an unidentified airborne pathogen. We showed the pathogen possessed the characteristics including airborne transmitted, cell-dependent, insensitive to antibiotics, filterable through 0.1 μm membrane. These properties prompt us to speculate that the pathogen of BSDs might be one kind of unidentified virus-like organism. However, the imperfection is that we have not isolated this putative pathogen from BSD+ samples [1]. Here, we report a further investigation of finding the virus-like pathogen in the serum and tissue extracts from BSD+ mice. Most importantly, this virus-like pathogen can be reisolated from the BSD+ extracts-inoculated, diseased BSD− cells.

To detect the putative virus-like pathogen in the BSD+ serum or tissue extracts, the serum and extracts of lung, liver, ovary, uterus, kidney were negatively stained and observed under TEM (transmission electron microscope). However, since excess impurities in the serum or tissue extracts caused overhigh background (electron beam were blocked), we had to dilute the samples 50 times before loading on the copper grids. The putative pathogen was captured and displayed a virus-like morphology (Fig. 1).

**Fig. 1.**
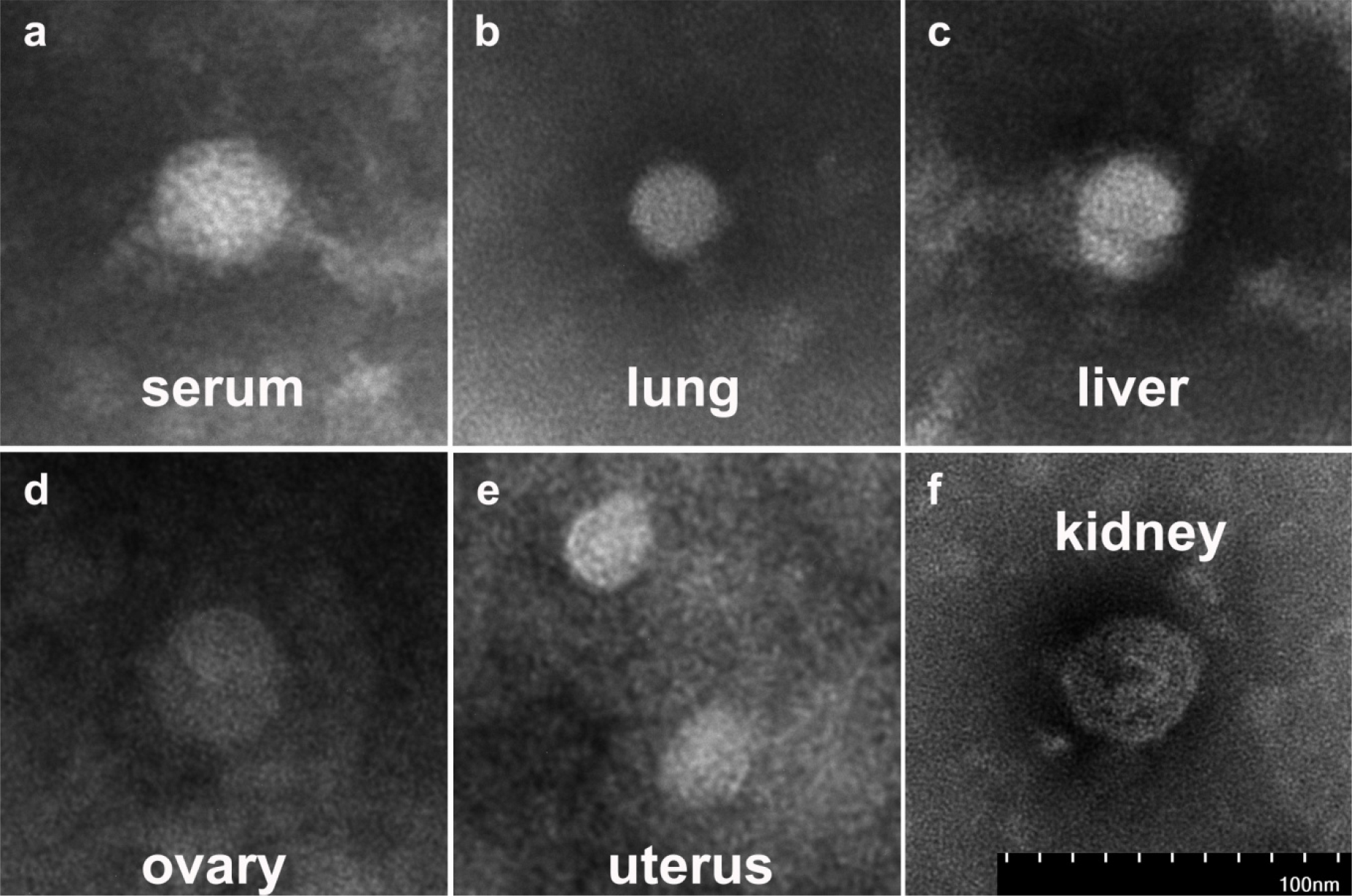
The representative image of the virus-like pathogen in the serum (a) and tissue extracts of lung (b), liver (c), ovary (d), uterus (e) and kidney (f) from BSD+ mice. The samples were diluted 50 times before loading on the copper grids to avoid overhigh background caused by excess impurities in the serum or tissue extracts. The samples were negatively stained with 2% phosphotungstic acid. Scale bar represents 100 nm.

The dilution made the pathogen very sparse under TEM. For concentration of the virus-like pathogen and elimination of the impurities, we collected all above mentioned tissues together and prepared the tissue extracts (BSD+ extracts). Then we performed ultracentrifugation and washed the precipitate with ddH_2_O. The resuspended precipitate was negatively stained and observed under TEM. The enriched virus-like pathogens were observed and mainly ranged from 20~40 nm (Fig. 2a). When BSD+ extracts were inoculated to BSD− cells, BSD− cells were transformed into BSD+ cells and began to produce BSDs, similar to our previous report [1]. Such a cytopathogenic effect (CPE) would appear in 1~3 days, which might depend on the numbers of the pathogen in different batch of extracts in the repeated experiments. In these transformed BSD− cells, the virus-like pathogens could be reisolated again (Fig. 2b).

**Fig. 2.**
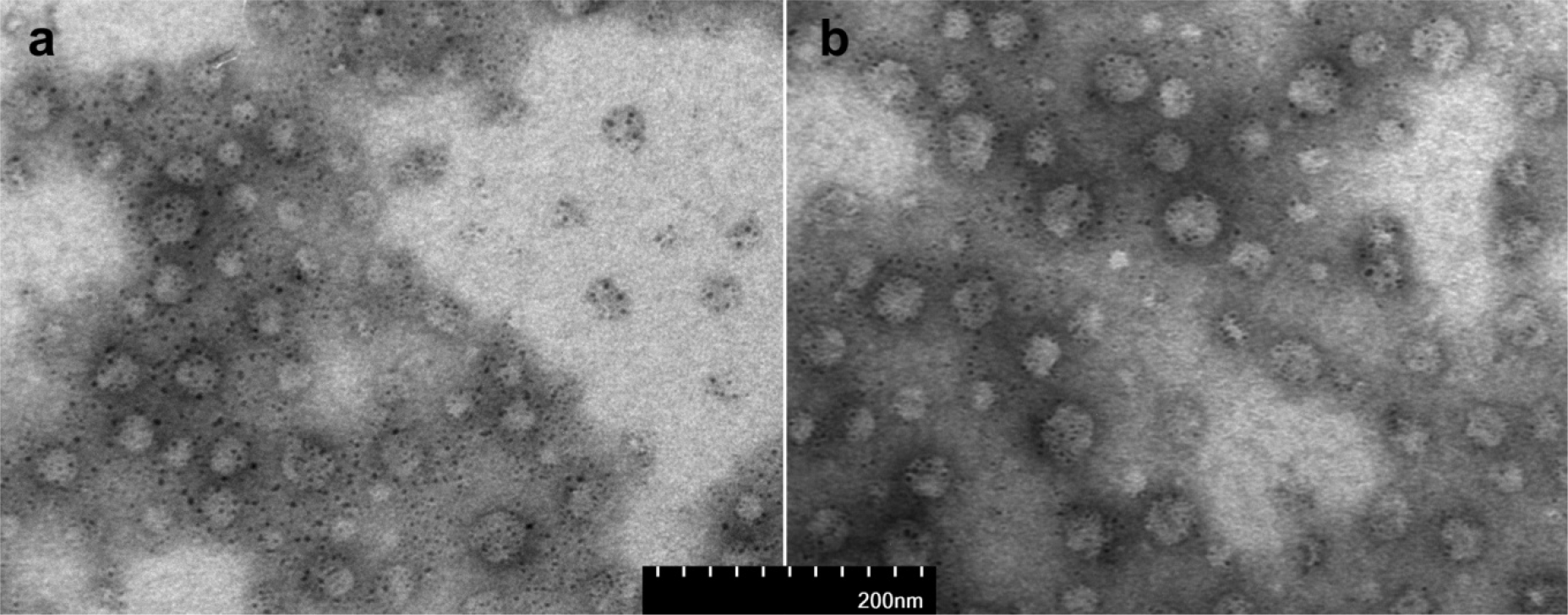
The representative image of the concentrated virus-like pathogen. The ultracentrifuged deposits were resuspended and negatively stained with 2% phosphotungstic acid. **a**, the sample isolated from BSD+ tissue extracts; **b**, the sample isolated from BSD+ extracts-inoculated, diseased BSD− cells. Scale bar represents 200 nm.

These findings support the speculation that BSDs are inorganic nanoparticles produced by virus-like pathogen infected cells. Furthermore, the findings here and the results in our previous study perfectly conform to Koch’s postulates. Therefore, we suggest that a kind of unidentified virus is the possible cause of BSDs in cell culture.

As we have mentioned in our previous paper, this research needs multidisciplinary cooperation [1]. We are now performing the nucleic acid isolation and sequencing. Nevertheless, we have limited knowledge of virus, so many virological protocols are new for us. Therefore, we provide these preliminary results and hope that, on the one hand, the independent laboratories test whether our results could be repeated; on the other hand, the professional virologists may sequence and classify the BSDs’ pathogen as quickly as possible.

## Methods

The methods were same as our previous paper [1], except the following.

### Preparation of extracts of different tissues

The lung, liver, ovary, uterus and kidney were dissected from BSD+ mice. The tissues were homogenized in DMEM/F12 media (about 0.1~0.5 g tissue/mL media), respectively. After 30 min on ice, the homogenates were centrifuged at 10,000 rpm for
15 min. The supernatants were filtered through 0.45 μm pore size membrane, then
0.22 μm membrane, and stored in 4 °C. Before inoculation into cell dishes or preparation for TEM, the extracts were centrifuged at 10,000 rpm for 5 min and filtered through a 0.1 μm pore size membrane. For TEM, the extracts were diluted 50 times to avoid overhigh background caused by excess impurities.

### Concentration of the virus-like pathogen and elimination of the impurities

Since the abover dilution made the pathogen very sparse, we performed ultracentrifugation for concentration of the pathogen and elimination of the impurities. The lung, liver, ovary, uterus and kidney from BSD+ mice were collected together and prepared extracts (BSD+ extracts) by using the above protocols. Before inoculation or TEM observation, BSD+ extracts were centrifuged at 10,000 rpm for 5 min and filtered through a 0.1 μm pore size membrane.

About 7 mL of the extracts were centrifuged at 10,000 rpm for 15 min. The supernatant was ultracentrifuged at 70,000 g for 40 min. The precipitate was resuspended and thoroughly washed with 7 mL of ddH_2_O, and then ultracentrifuged at 70,000 g for 40 min again. The precipitate was resuspended with 200 μL of ddH_2_O for TEM. For negative control, 7 mL of DMEM/F12 media were treated using the same protocols.

### BSD+ extracts inoculation of BSD cells and sample preparation for TEM

200~600 μL of BSD+ extracts were inoculated into the dishes of BSD− cells. Then, the CPE (BSDs appeared and multiplied) was monitored every day. For negative control, 200~600 μL of DMEM/F12 media were added into the dishes of BSD− cells. For empty control, 200~600 μL of BSD+ extracts were added into the dishes without cells. In negative control, CPE did not appear and no BSDs were observed. In empty control, even if BSD+ extracts were added, no BSDs grew out in one week.

For TEM, the BSD+ extracts-inoculated, diseased BSD− cells were freezed and thawed several times, and then 7 mL of ddH_2_O was added. The sample was centrifuged at 10,000 rpm for 15 min. The supernatant was ultracentrifuged at 70,000 g for 40 min. The precipitate was resuspended and thoroughly washed with 7 mL of ddH_2_O, and then ultracentrifuged at 70,000 g for 40 min again. The precipitate was resuspended with 200 μL of ddH_2_O for TEM. The negative control was prepared using the same protocols.

### Electron microscopy

About 25 μL of sample was placed directly on the BSA (bovine serum albumin) treated carbon-coated copper grids for 15 min. Then, the sample was absorbed with filter paper. After drying, the copper grids were stained with 2% phosphotungstic acid (pH 7.0), dried, and then examined under a Hitachi HT7700 TEM. The negative control showed no specific staining.

### Statistics

The investigators were not blinded during experiments. Each experiment was independently performed three times.

## Acknowledgements

We thank Dexian Zhang for assistance in animal management. This research was supported by the National Natural Science Foundation of China (Grant 31301201 to S.T.).

## Author Contributions

S.T. conceived the idea and designed the research; X.Y., X.J., W.A., S.W., Z.W & S.T. performed the experiments; X.L., H.L. & F.S. provided some materials and discussed the results; X.Y., X.J., W.A., H.L., S.M. & S.T. analyzed the data; S.T. wrote the manuscript.

